# Kidins220 promotes thymic iNKT cell development by reducing TCR signals, but enhances TCR signals in splenic iNKT cells

**DOI:** 10.1101/2022.04.05.487104

**Authors:** Laurenz A. Herr, Gina J. Fiala, Anna-Maria Schaffer, Katrin Raute, Rubí M.-H. Velasco Cárdenas, Jonas F. Hummel, Karolina Ebert, Yakup Tanriver, Susana Minguet, Wolfgang W. Schamel

**Affiliations:** Signaling Research Centers BIOSS and CIBSS; Department of Immunology, Faculty of Biology, University of Freiburg, Freiburg, Germany; Centre for Chronic Immunodeficiency (CCI), University of Freiburg, Freiburg, Germany, Faculty of Medicine; Spemann Graduate School of Biology and Medicine (SGBM), University of Freiburg, 79104, Freiburg, Germany; Institute of Medical Microbiology and Hygiene, Medical Center, University of Freiburg, Freiburg, Germany; Department of Medicine IV: Nephrology and Primary Care, Medical Center, University of Freiburg, Freiburg, Germany

**Keywords:** iNKT, Kidins220, ARMS, development, TCR, T cell

## Abstract

The stepwise development of thymic invariant natural killer T (iNKT) cells is controlled by the TCR signal strength. The scaffold protein Kinase D interacting substrate of 220 kDa (Kidins220) binds to the TCR regulating TCR signaling. T cell-specific Kidins220 knock-out (T-KO) mice contain severely decreased iNKT numbers. Very early in iNKT development TCR signals are reduced in the T-KO. In later steps, TCR signaling is increased in the T-KO leading to enhanced apoptosis of iNKT cells. Kidins220’s absence affects the iNKT1 subset most as it requires the weakest TCR signals for development. We also show that in iNKT1 development, weak TCR signals promote the progressive loss of CD4. In the periphery, Kidins220 switches its role back to promoting TCR signaling as splenic T-KO iNKT cells produce less cytokines and show reduced TCR signaling after *in vivo* stimulation with α-galactosylceramide. In conclusion, Kidins220 promotes or inhibits TCR signaling depending on the developmental context.

**summary statement:** We demonstrate that the transmembrane scaffold protein Kidins220 switches its role twice in iNKT cell biology: from a positive to a negative regulator of TCR signal strength during thymic development and back to a positive regulator in the periphery.

## Introduction

T lymphocytes are a crucial part of the adaptive immune system. αβ T cells, which express the αβ T cell antigen receptor (TCR) to recognize foreign and pathogenic antigens, can be further subdivided into conventional T cells, such as CD4^+^ T helper and CD8^+^ T killer cells, and unconventional T cells, such as invariant natural killer T (iNKT) cells.

iNKT cells are innate-like αβ T cells expressing a limited set of TCRα and TCRβ chains. Unlike conventional T cells, iNKT cells react on threats almost immediately. One of their main actions is the secretion of cytokines, such as interleukin 4 (IL-4), IL-17 and interferon-γ (IFN-γ) (Lee et al., 2013; Yoshimoto and Paul, 1994; Bendelac, 1995; Godfrey et al., 2000; Kronenberg and Gapin, 2002; Stetson et al., 2003; Crowe et al., 2003; Michel et al., 2007). Hence, they are crucial for initiating and regulating immune responses. With their invariant TCR they recognize a variety of glycolipids such as α-glucosyldiacylglycerol which originates from the cell wall of several bacteria, e.g., *S. pneumoniae*. These glycolipids are loaded onto and presented by the major histocompatibility complex I (MHC I)-like molecule CD1d on antigen presenting cells (APCs) (Kinjo et al., 2011).

All T cells are generated in the thymus and depending on their TCR’s specificity they are instructed to develop into conventional CD4^+^ or CD8^+^ T cells or among others into iNKT cells. To be positively selected to become conventional T cells, the TCR of CD4^+^ CD8^+^ (double positive, DP) thymocytes bind to peptide-loaded, classical MHC proteins present on thymic epithelial cells (Robey and Fowlkes, 1994; Robey et al., 1990; Kaye et al., 1989; Marusić-Galesić et al., 1989; Teh et al., 1988; Kruisbeek et al., 1985; Wang et al., 2020). To become positively selected to the iNKT lineage, DP thymocytes carrying the Vα14Jα18 TCR, bind to glycolipid-loaded CD1d molecules which are expressed on DP thymocytes (Bendelac et al., 1995; Coles and Raulet, 2000; Gapin et al., 2001). This binding generates a strong TCR signal that is required for iNKT cell selection (Moran et al., 2011). After selection, iNKT cells can develop along one of three developmental routes and thereby become iNKT1, iNKT2 or iNKT17 terminally differentiated iNKT cells (Lee et al., 2013). The developmental route taken depends on the iNKT cell’s TCR signal strength (Tuttle et al., 2018). Strong TCR signals result in the preferential generation of iNKT2 and iNKT17 cells. In contrast, weaker signals promote the development of iNKT1 cells. Thus, genetic mutations affecting positive regulators of TCR signaling and thereby reducing TCR signal strength, such as ZAP-70 (Hsu et al., 2009; Sakaguchi et al., 2003), were shown to decrease iNKT2 and iNKT17 cell numbers, whereas iNKT1 cell numbers were increased (Tuttle et al., 2018). Hence, iNKT2 and iNKT17 cells depend on strong TCR signals whereas iNKT1 cells rely on weaker signals.

In mice, iNKT2 cells express CD4 whereas iNKT17 cells do not. iNKT1 cells can either be CD4^+^ or CD4^-^ (Lee et al., 2013). Interestingly, the CD4^-^ and CD4^+^ iNKT1 subsets fulfil different functional roles with CD4^-^ iNKT1 cells possessing more NK cell-like functions such as tumor killing, whereas CD4^+^ iNKT1 cells express more proteins associated with helper T cell functions like IL-4 (Georgiev et al., 2016). The signals that drive CD4^+^ iNKT1 or CD4^-^ iNKT1 cell development remain unknown.

Directly after selection the developing iNKT cells are CD24^+^ CD44^-^ NK1.1^-^ and called stage 0 cells (iNKT0). They develop through stage 1 to stage 2 cells. Stage 1 (CD24^-^ CD44^-^ NK1.1^-^) already contains iNKT2 cells which are also found in stage 2 (CD24^-^ CD44^+^ NK1.1^-^). In stage 2 iNKT17 cells are also present. Stage 3 contains iNKT1 cells (CD24^-^ CD44^+^ NK1.1^+^) (Benlagha et al., 2002; Bennstein, 2018). Thus, in the thymus all three subsets (iNKT1, iNKT2, iNKT17) are present as mostly resident and terminally differentiated cells (Hogquist and Georgiev, 2020; Wang and Hogquist, 2018).

Since TCR signal strength directs iNKT cell development, it is important to gain more insight into how this strength is regulated. We previously identified the scaffold protein Protein Kinase D (PKD)-interacting substrate of 220 kDa (Kidins220), also called Ankyrin repeat-rich membrane spanning (ARMS), to bind to the TCR resulting in increased TCR signaling (Deswal et al., 2013). Kidins220 is a large protein with four transmembrane regions with both, N- and C-termini, reaching into the cytoplasm. It does not belong to the tetraspanin family and has various protein-protein interaction domains, such as 11 ankyrin repeats, a SAM domain, a proline-rich sequence and a PDZ-binding motif. Kidins220 is thought to act as a signaling hub (Neubrand et al., 2012; Good et al., 2011). It was initially discovered in neurons, where it interacts with the nerve growth factor receptor (NGFR) and Neurotrophin receptors like Trk (Iglesias et al., 2000; Chang et al., 2004; Arévalo et al., 2004).

In the adaptive immune system, Kidins220 binds to the TCR in T cells and the B cell antigen receptor in B cells. In both cases Kidins220 contributes to the activation of the MAP kinase/Erk pathway by binding to B-Raf (Deswal et al., 2013; Fiala et al., 2015). The function of Kidins220 was studied in a murine T cell line by utilizing an shRNA-based Kidins220 knock down (KD) (Deswal et al., 2013). Following TCR stimulation Erk and Ca^2+^ signaling were reduced in Kidins220 KD cells, although TCR levels on the surface were increased. This excluded the possibility of weaker signaling due to a reduced number of TCRs. Consequently, after TCR stimulation Kidins220 KD cells were less activated, as seen by reduced production of IL-2, IFN-γ and CD69. Hence, Kidins220 is an important positive regulator of TCR signaling *in vitro* (Deswal et al., 2013).

Here, we gained insight into the function of Kidins220 in T cells *in vivo*. To this end, we generated T cell-specific Kidins220 KO (T-KO) mice which showed strongly reduced iNKT cell numbers. Analyzing the T-KO iNKT cells, we found that in thymic development Kidins220 first enhances TCR signals at the stage 0 selection step of iNKT cells. Surprisingly, Kidins220 then switches its function to reduce TCR signal strength in stage 2 and 3 iNKT cells, thus preventing excessive apoptosis. In peripheral iNKT cells, Kidins220 again switches its role to enhance TCR-mediated signaling in response to antigen.

## Results

### Absence of Kidins220 reduces iNKT cell numbers

To study the role of Kidins220 in iNKT cell development, we generated a T cell-specific Kidins220 KO (T-KO) C57BL/6 mouse line. Mice with a floxed Kidins220 gene (Cesca et al., 2011, 2012) were crossed with mice expressing the Cre recombinase under the proximal Lck promotor (Orban et al., 1992; Hennet et al., 1995) which deletes floxed genes starting at the double negative 2 (DN2) stage of thymocyte development (Shimizu et al., 2001; Shi and Petrie, 2012; Fiala et al., 2019). T-KO mice carry the floxed Kidins220 gene on both alleles and LckCre, whereas control (Ctrl) mice carry at least one WT Kidins220 allele and LckCre (Fig. 1A). A flow cytometric analysis of the spleens shows that T cell numbers were diminished two-fold in T-KO compared to Ctrl mice, indicating that the T cell compartment is affected by the absence of Kidins220. As expected, B cell numbers were unaffected in T-KO mice (Fig. 1B).

**Figure 1.**
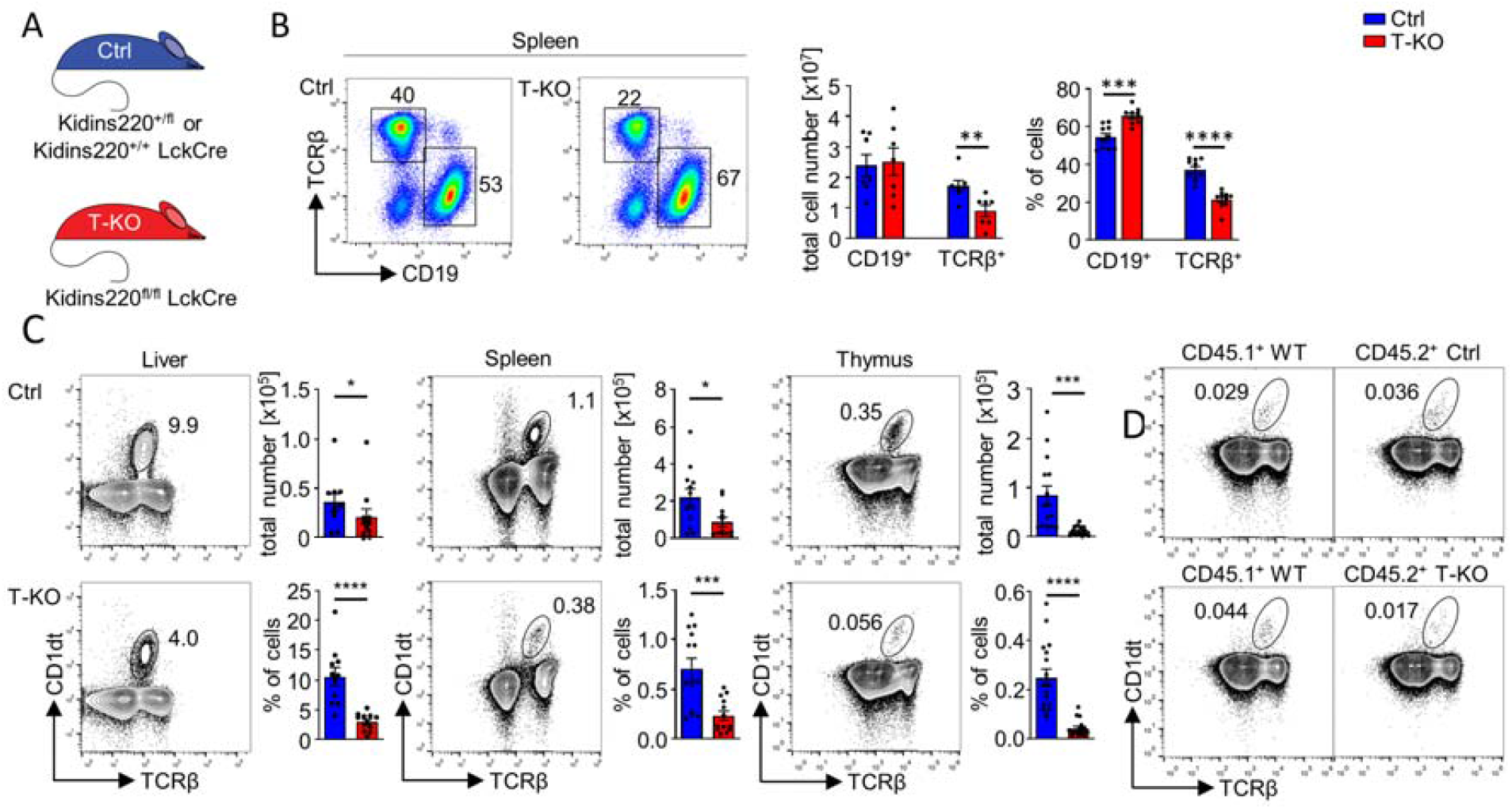
The numbers of conventional and un-conventional T cells are decreased in T-KO mice. (A) Schematics of the genotypes of Ctrl and T cell-specific Kidins220 KO (T-KO) mice are shown. (B) B and T cells in the spleen of Ctrl and T-KO mice were analyzed by anti-CD19 and anti-TCRβ staining. Total cell numbers and relative values are shown (n = 7-10). (C) iNKT cells were analyzed by staining lymphocytes from Ctrl and T-KO liver, spleen and thymus with CD1d tetramers (CD1dt) and anti-TCRβ antibodies. Total and relative cell numbers are depicted (n > 11). (D) Flow cytometric analyses of iNKT cells in the thymus of mixed bone-marrow chimeric mice are shown. RAG2 KO mice were lethally irradiated and reconstituted with bone marrow cells from CD45.1^+^ WT and CD45.2^+^ Ctrl or CD45.2^+^ T-KO mice in a 1:1 ratio. A representative analysis of the thymocytes of the WT/Ctrl (upper panel) and WT/T-KO (lower panel) chimeras is shown (n = 7-8). For this figure and all following ones, Ctrl samples are depicted in blue, and T-KO samples in red. Statistical analyses for total cell numbers was done by two-sided Student’s t test and for relative cell numbers by Mann-Whitney U test; * p < 0.05; ** p < 0.01; *** p < 0.001; **** < 0.0001. Error bars indicate SEM.

To test whether unconventional T cell numbers were also impacted by the loss of Kidins220, we analyzed iNKT cells by staining with CD1d-tetramers loaded with PBS57, an analog of α-Galactosylceramide (αGalCer). When bound to CD1d αGalCer (and PBS57) serve as potent ligands for the iNKT cell’s invariant TCR (Brossay et al., 1998; Kawano et al., 1997). In liver, spleen and thymus, we found a reduction of iNKT cell numbers in T-KO mice by 2-, 3- and 6-fold, respectively (Fig. 1C), suggesting that iNKT cell development is hampered in the absence of Kidins220.

During thymic development, iNKT precursor cells are selected by DP thymocytes presenting glycolipid-loaded CD1d molecules (Bendelac, 1995; Coles and Raulet, 2000). We found that CD1d levels were reduced by 1.2-fold in DP thymocytes of T-KO mice compared to Ctrl mice (Fig. S1A), while CD4 and CD8 levels showed no differences, indicating that there was not a general reduction in surface protein levels (Fig. S1B). The lower CD1d expression on DP cells could have caused a reduced selection of iNKT cells, since the latter need strong TCR signals for positive selection (Moran et al., 2011). To test whether the reduced CD1d expression could be the reason for inefficient iNKT cell development in T-KO mice, we generated bone marrow chimeras, for which equal amounts of bone marrow cells from CD45.1^+^ WT mice were mixed with CD45.2^+^ Ctrl or CD45.2^+^ T-KO bone marrow cells and injected intravenously into irradiated C57BL/6 Rag2^-/-^ (Rag2 KO) mice. In this setup, CD45.2^+^ T-KO iNKT precursor cells have access to normal amounts of CD1d molecules on the surface of CD45.1^+^ WT DP thymocytes. After eight weeks we analyzed iNKT cells in the recipient thymi. When WT cells were co-injected with Ctrl cells, the percentages of iNKT cells were almost the same in the WT CD45.1^+^ and Ctrl CD45.2^+^ populations (Fig. 1D, upper panel). However, after reconstitution with CD45.1^+^ WT and CD45.2^+^ T-KO bone marrow cells, the thymic T-KO iNKT cells showed a reduction of about 2-fold compared to the WT cells (Fig. 1D, lower panel and Fig. S1C). Hence, the decrease of iNKT cell numbers in T-KO mice was an iNKT intrinsic effect, since it could not be rescued once T-KO cells had access to sufficient amounts of CD1d during development.

### Kidins220 enhances T cell activation and TCR signaling in splenic iNKT cells

Next, we analyzed, whether Kidins220 plays a role for the execution of iNKT cell effector functions in the periphery. Since iNKT cells are a major source of early IL-4 and IFN-γ production after αGalCer challenge, we analyzed the production of these cytokines by splenic iNKT cells 2 hours after intraperitoneal (i.p.) αGalCer injection. By intracellular flow cytometric staining we found that the percentage of iNKT cells producing IL-4 or IFN-γ were reduced in T-KO compared to Ctrl mice (Fig. 2A and S2A). Further, the MFI for both cytokines was reduced in the cytokine-producing cells from T-KO mice. Injection of buffer alone did not lead to detectable cytokine production. Thus, T-KO iNKT cells were less activated than Ctrl iNKT cells upon TCR stimulation. This is in line with reduced TCR-mediated T cell activation in a conventional T cell line in which Kidins220 expression was downregulated (Deswal et al., 2013).

**Figure 2.**
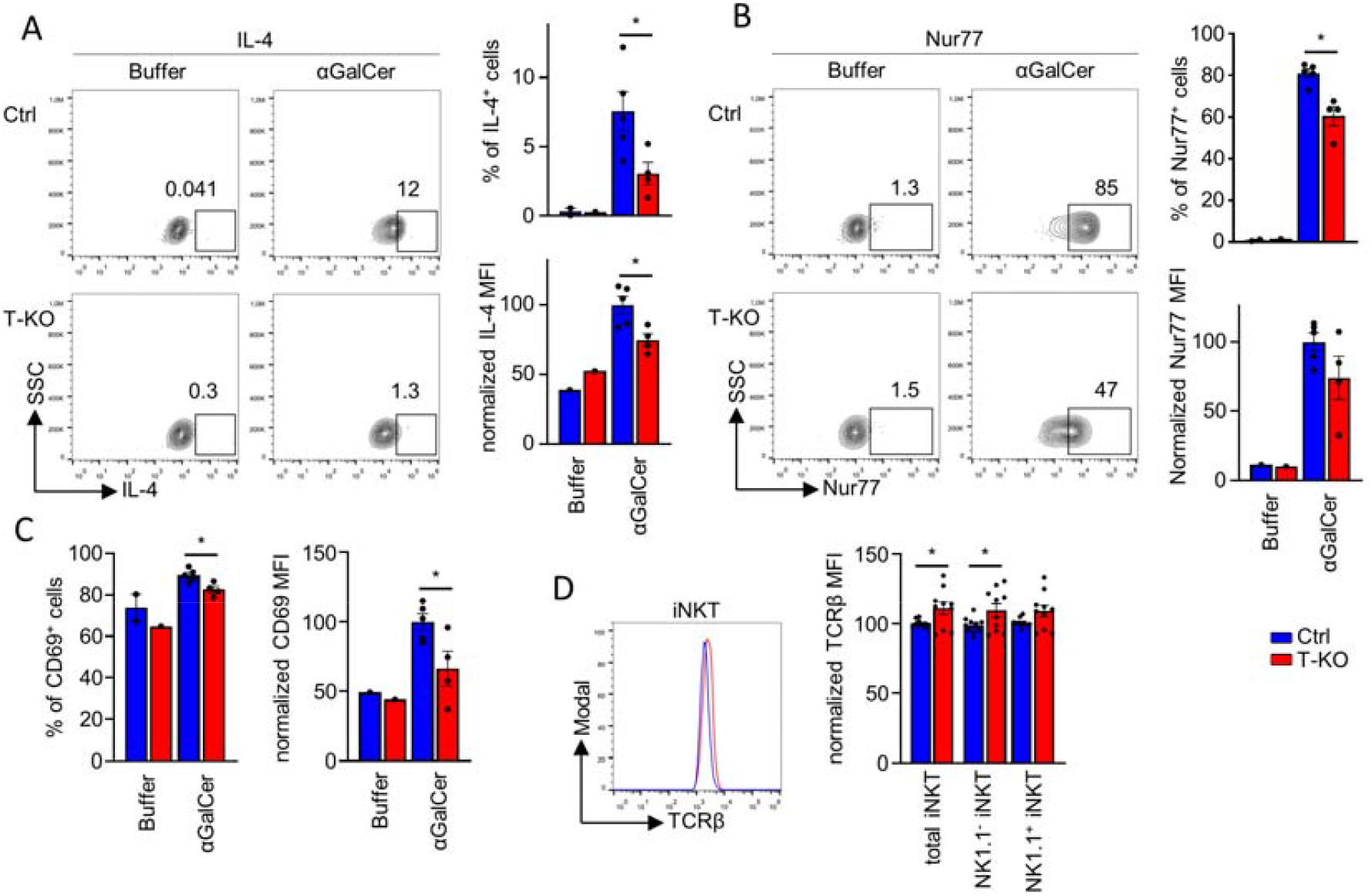
Reduced activation of iNKT cells from T-KO mice after αGalCer challenge. (A, B) Ctrl and T-KO mice were killed 2 h after i.p. injection of αGalCer or buffer alone. Expression of IL-4 (A) and Nur77 (B) was analyzed in iNKT cells by intracellular flow cytometry using anti-IL-4 and anti-Nur77 antibodies (n = 4-5). Plots were pre-gated on CD1dt^+^ TCRβ^+^ iNKT cells and MFI values were normalized to the ones of Ctrl samples for each independent experiment. (C) Statistics of the percent of CD69 expressing iNKT cells and of the CD69 MFI is shown after flow cytometry using anti-CD69 antibodies (n = 4-5). (D) TCRβ expression levels in total splenic iNKT cells or in iNKT cells, which were divided into NK1.1^-^ and NK1.1^+^ populations, were determined by flow cytometry (n = 10). Statistical analysis for relative cell numbers was done by Mann-Whitney U test and for MFIs by two-sided Student’s t test; * p < 0.05. Error bars indicate SEM.

Less production of IL-4 and IFN-γ could be due to reduced signaling via the iNKT cells’ TCR. Nur77 is expressed after TCR engagement, thus reporting on TCR signal strength (Moran et al., 2011; Cruz Tleugabulova et al., 2016; Kumar et al., 2020). We found that αGalCer-treated T-KO mice had 60%, whereas Ctrl mice had 81% Nur77^+^ iNKT cells in the spleen (Fig. 2B), and that the Nur77 expression levels in iNKT cells had the tendency of being lower in T-KO compared to Ctrl. This indicates that in splenic iNKT cells the absence of Kidins220 reduces TCR signaling. Indeed, diminished TCR signaling in T-KO iNKT cells would also explain the reduced CD69 expression levels following αGalCer challenge *in vivo* (Figs. 2C and S2B).

Splenic T-KO iNKT cells expressed 1.1-fold higher TCR levels compared to the Ctrl iNKT cells as detected by an anti-TCRβ stain. The same held true when separating total iNKT cells into NK1.1^-^ and NK1.1^+^ iNKT cells (Fig. 2D). This is consistent with findings that a downregulation of Kidins220 in a mature conventional T cell line led to slightly increased TCR levels (Deswal et al., 2013), and excludes that less TCR signaling in the absence of Kidins220 was due to lower TCR levels.

In conclusion, Kidins220 promotes TCR signaling in peripheral iNKT cells. This is in line with its binding to the TCR and the kinase B-Raf in a conventional T cell line (Deswal et al., 2013).

### Development of T-KO iNKT cells is impeded in T-KO mice, with strongly reduced iNKT1 numbers

Reduced iNKT cell numbers in T-KO mice might originate from an inefficient development of these cells in the thymus. To elucidate iNKT cell development, we analyzed thymic iNKT cells (CD1dt^+^ TCRβ^+^) from Ctrl and T-KO mice by flow cytometry. These cells were divided into the established stages 0 and 1 (stage 0 + 1, CD44^-^ NK1.1^-^), stage 2 (CD44^+^ NK1.1^-^) and stage 3 (CD44^+^ NK1.1^+^; Fig. 3A). Further, stage 0 was assessed separately (CD44^-^ CD24^+^; Fig. 3B). Similar total iNKT cell numbers in stage 0 + 1 and in stage 0 alone were detected in T-KO and Ctrl mice. In sharp contrast, stage 2 and 3 cell numbers were reduced in T-KO mice, with stage 3 cells being affected the most (Fig. 3A). This reduction led to a percentwise increase of cells in stages 0 and 1 and a decrease in stage 3. Furthermore, CCR7^+^ iNKT precursor cells (Wang and Hogquist, 2018) were equal in numbers in Ctrl and T-KO mice (Fig. 3C), being in line with equal stage 0 numbers.

**Figure 3.**
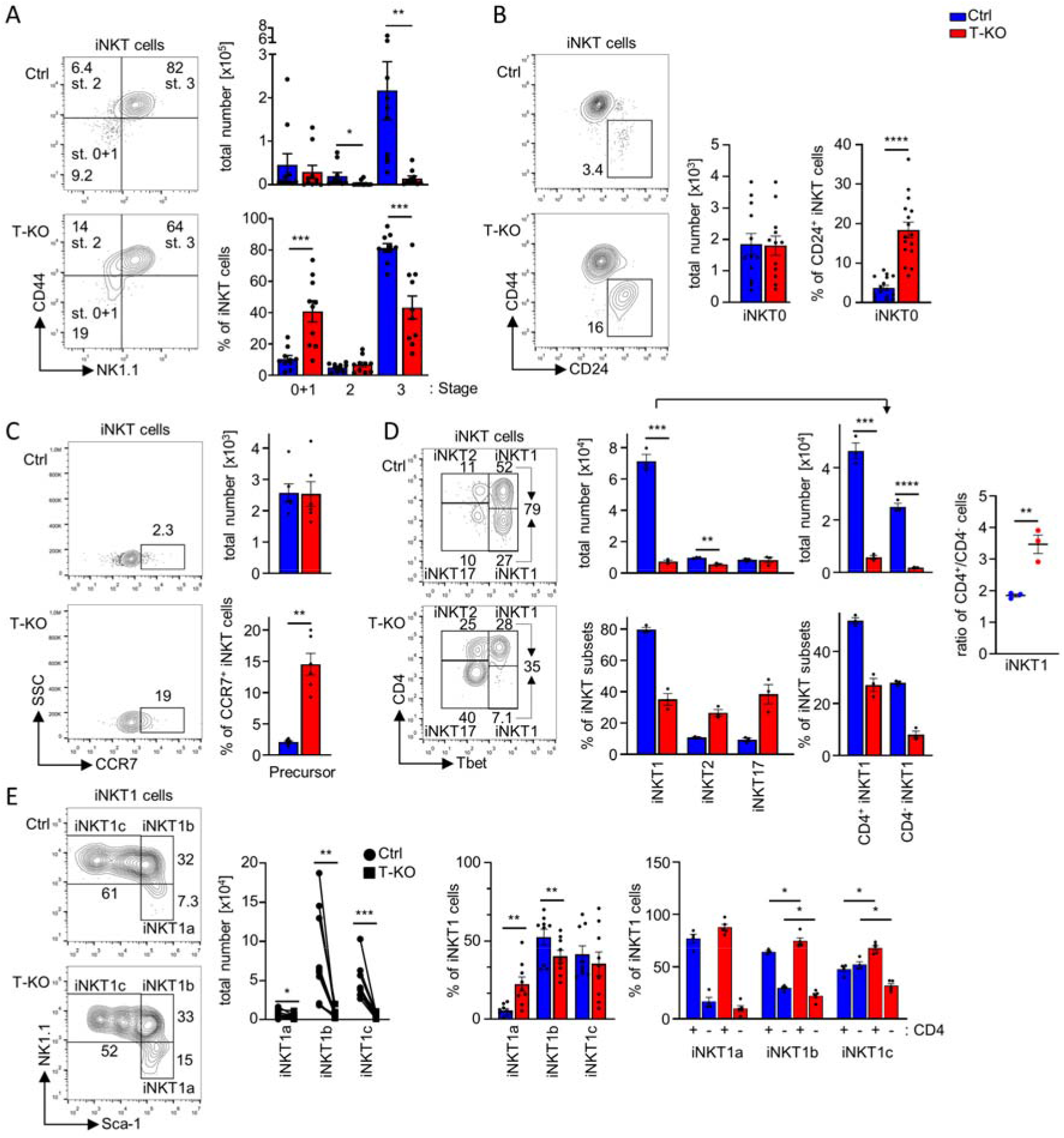
Partial block of iNKT development from stage 2 to stage 3 in T-KO mice. (A) CD1dt^+^ TCRβ^+^ iNKT cells were analyzed by flow cytometry using anti-CD44 and anti-NK1.1 antibodies and grouped into stages 0 + 1, 2 and 3. Graphs show total iNKT and relative cell numbers (n = 10). (B) The amount of CD24^+^ iNKT0 cells was analyzed using anti-CD24 and anti-CD44 antibodies (n = 12-16). (C) The expression of CCR7^+^ identifying iNKT precursor cells is depicted (n = 6). (D) Thymocytes were stained with anti-CD4 and with anti-Tbet antibodies intranuclearly (n = 3). (E) iNKT1 cells were subdivided into iNKT1a, b and c cells by using anti-Sca-1 and anti-NK1.1 antibodies (n = 11). Subsequently, each subset was divided into CD4^+^ and CD4^-^ populations using anti-CD4 antibodies (n = 4). In all panels, cells were pre-gated for CD1dt^+^ TCRβ^+^ iNKT cells. Statistical analysis for relative cell numbers was performed by Mann-Whitney U test, and for total cell numbers by two-sided Student’s t test and for paired analysis of total cell numbers by Wilcoxon matched-pairs signed rank test; * p < 0.05; ** p < 0.01; *** p < 0.001; **** < 0.0001. Error bars indicate SEM.

The strong reduction of stage 3 iNKT cell numbers suggested that iNKT1 cell numbers were reduced. The iNKT1 cells express the transcription factor Tbet and are either positive or negative for CD4. The iNKT2 cells are Tbet^-^ CD4^+^, whereas iNKT17 cells are Tbet^-^ CD4^-^ (Lee et al., 2013). Indeed, numbers of iNKT1 cells were reduced in T-KO mice 10-fold, iNKT2 cell numbers 2-fold and iNKT17 cell numbers were unchanged (Fig. 3D). This shows that the absence of Kidins220 strongly interferes with iNKT1, slightly with iNKT2 and not with iNKT17 development. Numbers of both CD4^+^ and CD4^-^ iNKT1 cells were reduced; with CD4^-^ iNKT1 cells being affected stronger (Fig. 3D). Using CD122 as an alternative strategy to identify iNKT1 cells (Georgiev et al, 2016), we also saw a strong reduction of these cells in the absence of Kidins220 (Fig. S3).

Since iNKT1 (NK1.1^+^ CD44^+^) cells showed the strongest reduction, we subdivided those cells into iNKT1a, b and c subsets based on Sca-1 and NK1.1 expression (Baranek et al., 2020). iNKT1a cell numbers were only slightly reduced in T-KO mice. (Fig. 3E). However, there was a strong reduction of iNKT1b and iNKT1c cell numbers, indicating that the partial developmental block might occur at the transition from iNKT1a to iNKT1b in T-KO mice.

Since the ratio of CD4^+^ to CD4^-^ iNKT1 cells was larger in T-KO mice (Fig. 3D), we tested at which stage the CD4^-^ iNKT1 cells would be reduced. In Ctrl mice, there were 4 times and 2 times more CD4^+^ than CD4^-^ iNKT1a and iNKT1b cells, respectively (Fig. 3E). This was unchanged in the T-KO mice. In Ctrl mice CD4^-^ cells seem to “catch up” at the iNKT1c cell subset, since there were equal numbers of CD4^+^ and CD4^-^ cells. However, in T-KO mice there were still more CD4^+^ than CD4^-^ iNKT1c cells. Thus, the loss of Kidins220 interferes with the later stages of iNKT1 development, and hinders CD4^-^ iNKT1c cells to develop.

### T-KO iNKT cells are more susceptible to apoptosis and show higher proliferative activity

In the absence of Kidins220, iNKT cell numbers were increasingly reduced the further they progressed in development. Hence, we tested whether iNKT T-KO cells exhibit increased apoptosis. To this end, thymocytes were cultured overnight *ex vivo* and subsequently stained to visualize active Caspase 3/7 by flow cytometry. In thymi of T-KO mice about 4 times more iNKT cells were apoptotic compared to Ctrl mice (Fig. 4A), which was further confirmed by Annexin V staining (Fig. S4A). As seen by 7-AAD staining, thymic T-KO iNKT cells contained higher percentages of dead cells than the ones of Ctrl mice (Fig. S4B). In line with the strongest reduction of NK1.1^+^ stage 3 iNKT cell numbers in T-KO mice (Fig. 3A), the increase of apoptosis (T-KO vs Ctrl) was stronger in NK1.1^+^ iNKT cells compared to the one of NK1.1^-^ iNKT cells (Fig. 4B). In general, apoptosis was more prominent in NK1.1^-^ iNKT cells compared to NK1.1^+^ iNKT cells and this is in line with previous studies (Lu et al., 2019).

**Figure 4.**
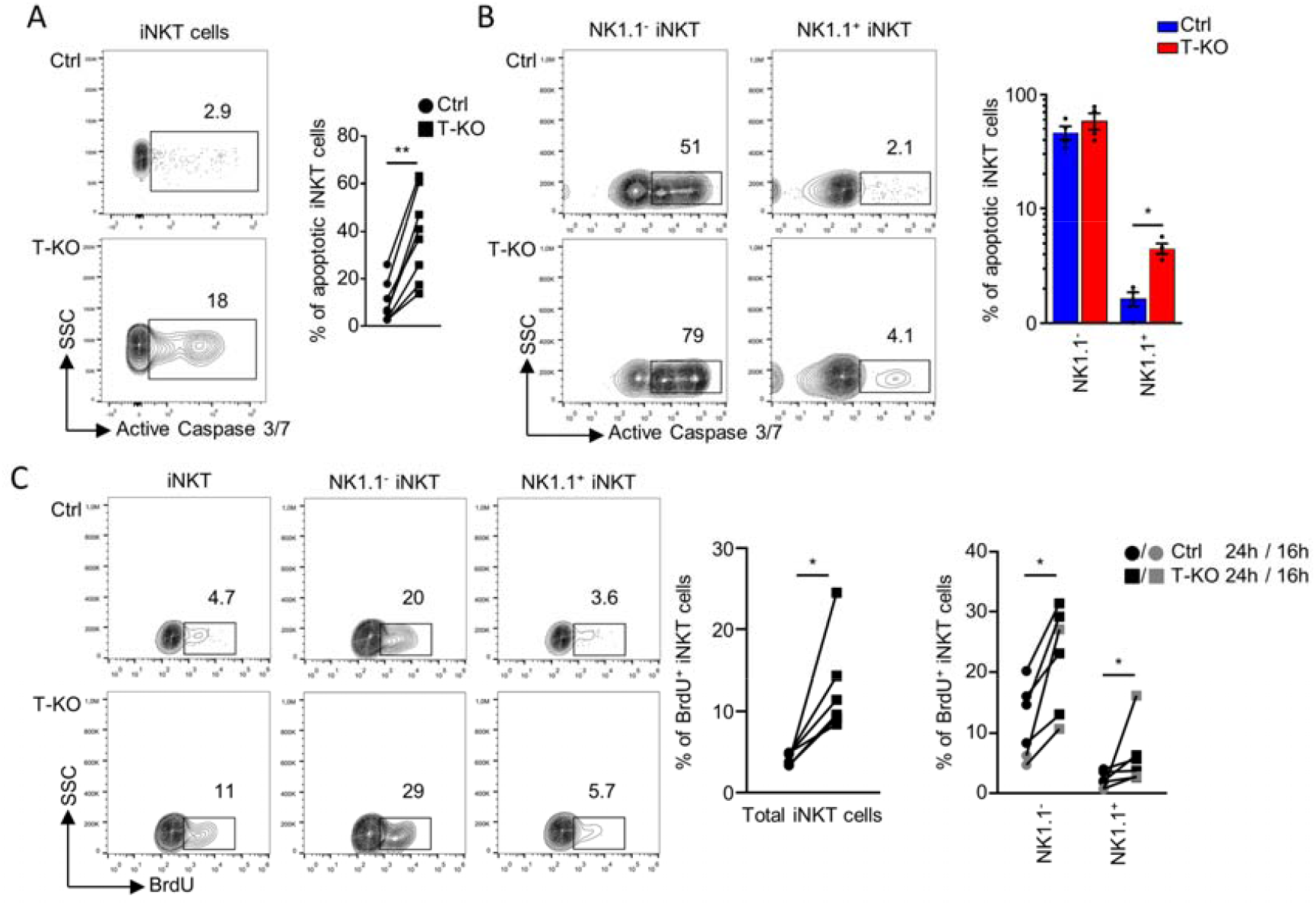
iNKT cells of T-KO mice are more proliferative and more susceptible to apoptosis compared to Ctrl cells. (A) To quantify apoptosis in iNKT cells, total thymocytes were cultivated for 18 h in RPMI medium supplemented with 10% FBS. Graph shows percentages of CD1dt^+^ TCRβ^+^ iNKT cells with active Caspase 3/7. Dead cells were excluded based on FSC and SSC values (n = 8). (B) Apoptotic iNKT cells as in (A) were subdivided into NK1.1^-^ and NK1.1^+^ cells (n = 4). (C) Ctrl and T-KO mice were i.p. injected with BrdU to analyze proliferating cells. BrdU-treated mice were killed 16 (grey) or 24 h (black) after BrdU injection and thymocytes were stained with anti-BrdU antibodies. CD1dt^+^ TCRβ^+^ iNKT cells and iNKT cells which were subdivided into NK1.1^+^ and NK1.1^-^ iNKT cells were analyzed using flow cytometry. Graphs show relative numbers of BrdU^+^ cells (n = 6-7). Statistical analysis for relative cell numbers in (B) was done by Mann-Whitney U test and in (A) and (C) by Wilcoxon matched-pairs signed rank test; * p < 0.05; ** p < 0.01. Error bars indicate SEM.

The anti-apoptotic protein BCL2 was expressed to the same levels in Ctrl and T-KO iNKT cells (Fig. S4C), suggesting that reduced BCL2 was not the cause for enhanced apoptosis in T-KO iNKT cells. Indeed, besides BCL2 (Yao et al., 2009), BCL_XL_ was also shown to be important for cell survival during iNKT cell development (Egawa et al., 2005), and this might have been altered in the T-KO cells.

To test for iNKT cell proliferation *in vivo*, we injected BrdU i.p. into T-KO and Ctrl mice. Since BrdU incorporates into the DNA during DNA synthesis, the proliferated cells can be visualized by flow cytometry using anti-BrdU antibodies. Unexpectedly, T-KO iNKT cells had proliferated more compared to the Ctrl cells (Fig. 4C). This held true for NK1.1^-^ and NK1.1^+^ iNKT cells.

In conclusion, Kidins220-deficient thymic iNKT cells proliferated more, but also were more prone to apoptosis than the Ctrl cells. And the latter might explain the reduction of iNKT cell numbers in T-KO mice.

### TCR signaling is reduced in stage 0 thymic T-KO iNKT cells and elevated in stage 2 and 3 T-KO iNKT cells

Enhanced proliferation and apoptosis in thymic T-KO iNKT cells could be caused by stronger TCR signaling during development and this can be measured by CD5 surface expression that correlates with the TCR signal strength (Azzam et al., 1998, 2001). Indeed, in total iNKT cells from T-KO mice we found elevated CD5 levels of about 1.4-fold compared to the ones from Ctrl mice (Fig. 5A). Interestingly, CD5 expression levels on the different iNKT stages is differentially affected by the absence of Kidins220: at stage 0 the CD5 levels were reduced in the T-KO, at stage 1 the levels of Ctrl and T-KO cells were similar and at stages 2 and 3 the CD5 levels were increased in the T-KO cells. Elevated CD5 expression was preserved in peripheral NK1.1^+^ and NK1.1^-^ iNKT cells from the spleen (Fig. 5B).

**Figure 5.**
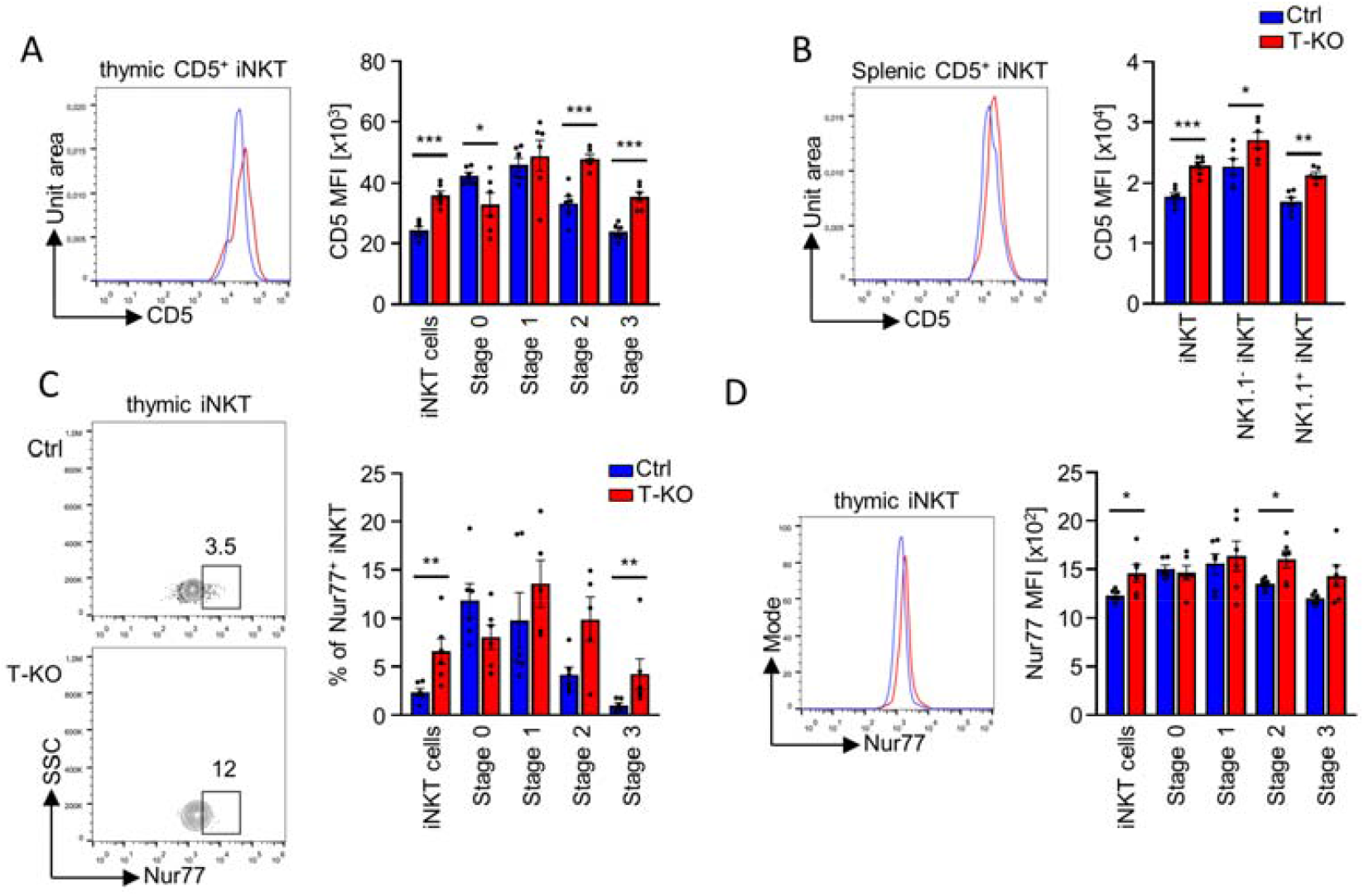
Stage 2 and 3 iNKT cells of T-KO mice exhibit stronger TCR signaling compared to Ctrl cells. (A) CD5 MFI in thymic iNKT cells was assessed by flow cytometry using anti-CD5 antibodies. Cells were pre-gated on CD1dt^+^ TCRβ^+^ iNKT cells. iNKT stages were analyzed using anti-CD24, anti-CD44, and anti-NK1.1 antibodies (n = 6). (B) CD5 MFI was assessed in splenic iNKT cells divided into NK1.1^+^ and NK1.1^-^ iNKT cells using anti-NK1.1 antibodies (n = 6). (C) Total thymic iNKT cells and iNKT stages as distinguished in (A) were analyzed regarding Nur77^+^ cells using anti-Nur77 antibodies. Graphs show relative numbers of Nur77^+^ iNKT cells (n = 6). (D) Nur77 MFI in iNKT cells as in (A) was determined by flow cytometric analysis using anti-Nur77 antibodies (n = 6). Statistical analysis for relative cell numbers was determined by Mann-Whitney U test and for MFIs by two-sided Student’s t test; * p < 0.05; ** p < 0.01; *** p < 0.001. Error bars indicate SEM.

As Nur77 expression levels can also be used to quantify TCR signal strength (Moran et al., 2011), we analyzed the percentage of Nur77^+^ iNKT cells in the thymus. In the total iNKT population there were three times more Nur77^+^ iNKT cells in T-KO mice compared to Ctrl mice (Fig. 5C). Consolidating the CD5 data, we again detected a lower (stage 0), equal (stage 1) and higher (stages 2 and 3) percentage of Nur77^+^ T-KO cells compared to Ctrl cells (Fig. 5C). In stage 3 the largest increase of Nur77^+^ in the T-KO cells was found (namely 4-fold compared to Ctrl). Nur77 expression levels in the cells as quantified by the MFI revealed the same trend (Fig. 5D).

Strikingly, these data suggest that Kidins220 enhances TCR signaling in thymic stage 0 iNKT cells but reduces TCR signaling at stages 2 and 3.

This change in TCR signaling might be due to altered TCR expression levels. Indeed, in total iNKT cells we detected increased TCR levels in the T-KO cells (Fig. 6A). In stage 0 TCR levels were lower, in stage 1 equal and in stages 2 and 3 higher in the T-KO compared to the Ctrl. The higher levels in stages 2 and 3 are in line with increased TCR levels of T-KO iNKT cells in the spleen (Fig. 2D) and in a T cell line where Kidins220 expression was downregulated (Deswal et al., 2013).

**Figure 6.**
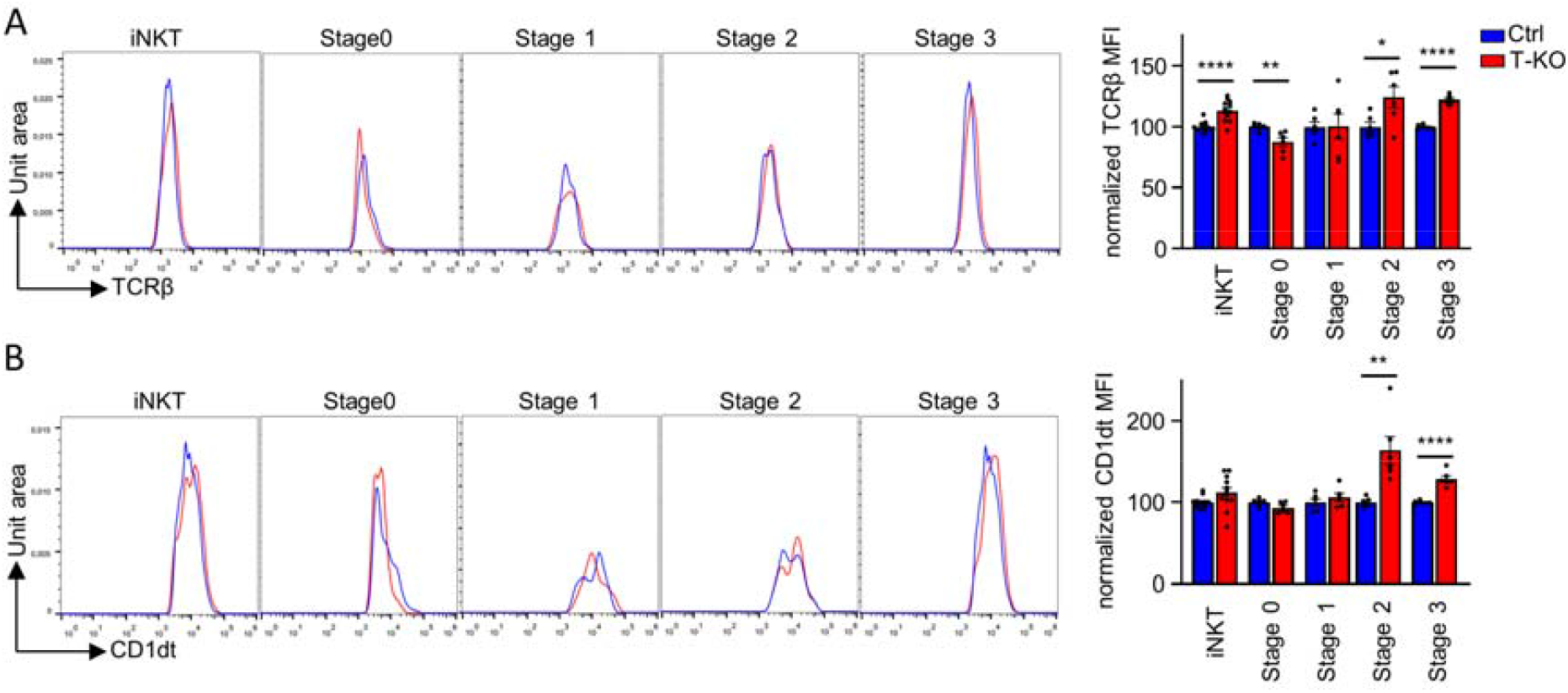
In T-KO mice stage 2 and 3 iNKT cells have increased TCR levels. (A) TCRβ expression levels and (B) CD1dt binding were analyzed by flow cytometry in iNKT cells. Cells were pre-gated on CD1dt^+^ TCRβ^+^ iNKT cells. iNKT stages were analyzed using anti-CD24 and anti-CD44 antibodies to visualize stage 0 iNKT cells and anti-CD44 and anti-NK1.1 antibodies were utilized to stain stages 1, 2, and 3. MFIs are depicted for (A) and (B) (n = 6-12). Statistical analysis for MFIs was performed by two-sided Student’s t test; * p < 0.05; ** p < 0.01; **** < 0.0001. Error bars indicate SEM.

Increased TCR levels correlated with increased binding of the TCR’s ligand CD1dt to the iNKT cells (Fig. 6B). Since more surface TCR on iNKT cells might lead to more TCR signaling (Tuttle et al., 2018), the enhanced CD1d binding we detected in the T-KO stage 2 and 3 cells might contribute to the stronger TCR signals observed in Figure 5, causing more apoptosis in those iNKT cells (Fig. 4).

## Discussion

Here, we show that the T cell-specific KO of Kidins220 leads to a severe reduction of iNKT cell numbers, whereas the number of conventional T cells is only slightly reduced. Surprisingly, Kidins220 switches its role twice in iNKT cell biology: from promoting TCR signaling in the selection of DP thymocytes to the iNKT lineage, to inhibiting signaling during iNKT cell development limiting TCR signal strength and allowing iNKT1 cells to develop, and back to promoting TCR signaling in peripheral iNKT cells (Fig. 7). During the development of iNKT cells, the TCR signal strength progressively decreases from developmental stage 0 to stage 3 (Lu et al., 2019; Moran et al., 2011). Like others (Moran et al., 2011; Azzam et al., 1998; Lu et al., 2019; Cruz Tleugabulova et al., 2016), we have used CD5 and Nur77 expression as readouts for TCR signal strength to evaluate the role of Kidins220 in iNKT cell development. In stage 0, the TCR signal is lower in T-KO cells compared to Ctrl, suggesting that Kidins220 is a positive regulator of TCR signaling (Fig. 7). This is in line with data derived from a murine T cell line in which TCR signaling was reduced when Kidins220 was knocked down (Deswal et al., 2013), and from cardiovascular and neurological systems, where Kidins220 positively couples several receptors to intracellular signaling (Arévalo et al., 2004). However, although strong signals are required for iNKT positive selection (Dutta et al., 2013; Seiler et al., 2012; Moran et al., 2011), we did not find reduced cell numbers in stage 0 in T-KO mice. Most likely, the lower signals were still above a threshold required for the DP cells to be selected to the iNKT lineage. TCR signal strength also steers development of the selected iNKT cells into the different subsets (iNKT1, iNKT2 and iNKT17). iNKT1 cells, which require a weak TCR signal to develop (Tuttle et al., 2018), were affected most by the T-KO being reduced 10-fold. Indeed, remaining iNKT1 cells exhibited increased TCR signaling (CD5 and Nur77) (stage 3 cells are mostly iNKT1 cells) and enhanced apoptosis. We also found increased proliferation in stage 3 iNKT cells (iNKT1), but this obviously could not compensate for the enhanced apoptosis. The generation of iNKT17 cells is promoted by TCR signals of medium strength and indeed their cell numbers were similar in T-KO and Ctrl. The generation of iNKT2 cells benefits from strong TCR signals and was slightly diminished (2-fold). As we did not see elevated apoptosis of iNKT2 cells, we expected an increase in the number of those cells in T-KO mice. In support of our finding, it was already shown that stronger TCR signaling in iNKT cells leads to a slight reduction of iNKT2 cell numbers (Lu et al., 2019).

**Figure 7.**
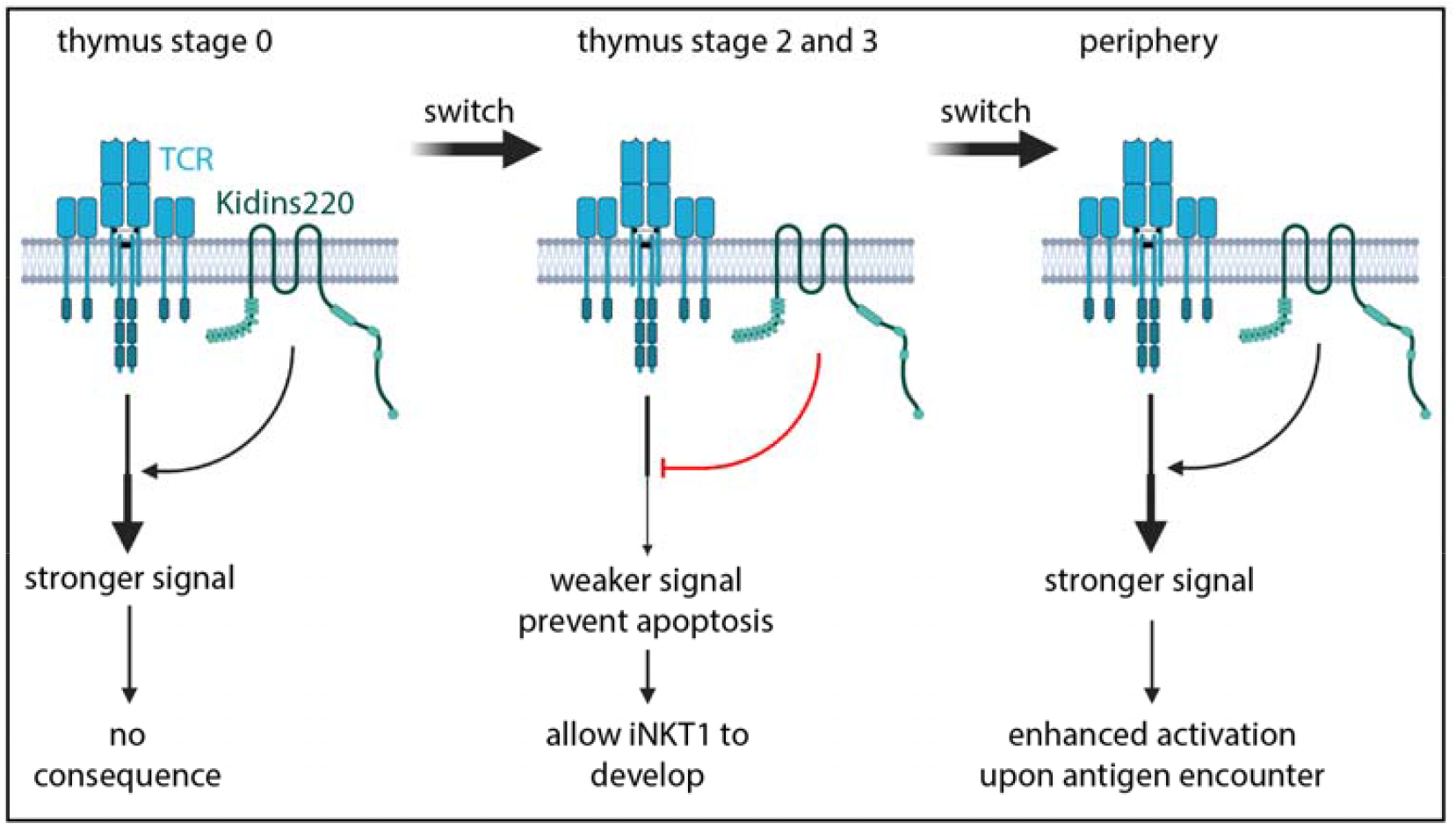
Kidins220 switches its function from a positive, to a negative and back to a positive TCR signal regulator during iNKT cell development and function. During stage 0 in thymic development, Kidins220 leads to stronger TCR signaling in iNKT cells. During stages 2 and 3 of development, this function switches to the opposite as Kidins220 dampens TCR signaling. In the periphery, after stimulation with αGalCer, Kidins220 again switches its role by enhancing TCR signaling. Created with BioRender.com.

Interestingly, Kidins220 shows progressively higher expression levels from stage 1 to stage 3 (https://www.immgen.org/), supporting the assumption that in later stages of development Kidins220 is needed the most to dampen TCR signals. This would make iNKT1 cells, which are mostly stage 3 cells, the most Kidins220-dependent iNKT subtype – as we found. Conversely, in the hypomorphic ZAP-70 mice, TCR signaling was reduced in stages 2 and 3, and iNKT1 cells were percentwise increased whereas iNKT2 and iNKT17 cells were reduced (Hsu et al., 2009; Sakaguchi et al., 2003; Tuttle et al., 2018; Zhao et al., 2018).

The phenotype of T-KO mice is very similar to the Slam family receptors (SFRs)-deficient mice (Lu et al., 2019), suggesting that these molecules might play similar roles in iNKT cells. For example, stage 3 iNKT cell numbers were diminished the most in SFR KO mice and TCR signaling was also elevated in stage 2 and 3 iNKT cells. However, there are differences to the Kidins220 KO, such as that Nur77 was equal in stage 0 SFR KO compared to the Ctrl, whereas in T-KO mice it was lower. Moreover, BCL2 expression was lower in SFR KO iNKT cells, explaining the enhanced apoptosis, whereas in iNKT cells lacking Kidins220 BCL2 levels remained unchanged.

CD1d, the ligand for the iNKT TCR, is expressed at slightly lower levels in the T-KO mice. Using bone marrow chimeric mice in which T-KO iNKT precursors have access to sufficient amounts of CD1d, we showed that iNKT cell numbers were also diminished, substantiating our conclusion that it is an iNKT cell intrinsic mechanism that led to reduced iNKT cell numbers, namely an altered TCR signal strength caused by the absence of Kidins220. Indeed, enhanced TCR signaling in the T-KO developing iNKT cells cannot be explained by lower CD1d levels.

In conclusion, the function of Kidins220 switches during development. In stage 0 Kidins220 promotes TCR signaling, and in stages 2 and 3 it dampens TCR signaling (Fig. 7). This is specific to Kidins220, since ZAP-70 was a positive regulator at all stages (Tuttle et al., 2018). Intriguingly, in peripheral iNKT cells Kidins220 switched back to its positive regulatory function. In fact, TCR signaling as measured by Nur77 expression was reduced after stimulation with αGalCer in splenic T-KO iNKT cells (Fig. 7). This resulted in lower iNKT cell activation as expression of CD69 and IL-4 was reduced. This is in line with Kidins220’s positive function in stage 0 cells and in a murine T cell line (Deswal et al., 2013).

How is the changing role of Kidins220 on TCR signaling regulated? This might be at two levels; firstly, by controlling TCR expression levels and secondly, by coupling to different signaling proteins.

Firstly, Kidins220 is involved in regulating receptor levels on the cell surface. When Kidins220 was downregulated in neurons or in a T cell line, glutamate receptor 1 or TCR levels were increased, respectively (Arévalo et al., 2010; Deswal et al., 2013). Likewise, TCR levels were increased in T-KO compared to Ctrl in stage 2 and 3 thymic and in splenic iNKT cells. Since in in stage 2 and 3 iNKT cells TCR signals were enhanced, but reduced in splenic iNKT cells, Kidins220 has other means to regulate signaling besides controlling receptor levels (see below). However, in all thymic iNKT cell stages TCR signal strength correlated with the TCR surface levels in T-KO versus Ctrl: in stage 0 cells TCR levels and signaling were reduced but increased in stage 2 and 3 cells in the T-KO and in stage 1 cells TCR levels and signaling were the same in T-KO and Ctrl.

Secondly, a different role of Kidins200 in the different iNKT cell developmental stages might also be accomplished through either binding to a protein that promotes signaling (in precursor and peripheral iNKT cells) or that reduces signaling (in thymic stage 2 and 3 iNKT cells). In a T cell line, it was shown that Kidins220 binds to the TCR and to B-Raf and thus connects the TCR to the MAP kinase/Erk pathway and couples the TCR to calcium signaling (Deswal et al., 2013). Binding to and coupling receptors to these pathways is a general function of Kidins220, as it was also demonstrated in B cells and neurons (Fiala et al., 2015; Arévalo et al., 2004; Jaudon et al., 2021; Cesca et al., 2011; Scholz-Starke and Cesca, 2016). Indeed, Ras, being part of the Erk pathway, is involved in iNKT development, since expression of a dominant negative Ras led to reduced iNKT cell numbers (Hu et al., 2011). The function of Kidins220 to couple the TCR to calcium influx is in line with the fact that Nur77 transcription relies not only on TCR-mediated signals, but that calcium signaling was the most important player (Liu, 2009; Youn Hong-Duk, Chatila Talal A., 2000; Lith et al., 2020). Kidins220’s putative binding to a negative regulator of TCR signaling in developing iNKT cells remains speculative. However, a changing role of Kidins220 was also shown in the nervous system when triggering TrkB. In embryonic astrocytes, Kidins220 promotes kinase-based signaling, and after birth Ca^2+^-dependent signaling (Jaudon et al., 2021).

Finally, our data shed some light on the generation of CD4^-^ and CD4^+^ iNKT1 cells. iNKT1a cells which are mostly CD4^+^ differentiate stepwise to iNKT1b and 1c. We show here that in these steps CD4 is progressively lost, until the number of CD4^+^ and CD4^-^ cells is similar in the iNKT1c subset. Interestingly, total cell numbers of iNKT1a cells are similar and the ratio of CD4^+^ and CD4^-^ iNKT cells are equal in T-KO and Ctrl, showing that Kidins220 has little effect during this developmental decision. Only in the following two stages, iNKT1b and 1c, T-KO iNKT1 cell numbers are decreasing compared to Ctrl, with CD4^-^ being reduced more than CD4^+^ cells, showing that Kidins220 promotes the generation of CD4^-^ iNKT cells. This suggests that the TCR signal strength also plays a role in the decision of whether an iNKT1 cells loses or preserves CD4 expression, in that weak signals promote the generation of CD4^-^ iNKT1 cells.

Reduced TCR signaling in stage 0 and enhanced signaling in later stages of T-KO iNKT cells, are a further support for the notion that iNKT cells receive persisting TCR engagement also at later stages (Park et al., 2019; Hogquist and Georgiev, 2020).

In conclusion, we show that the scaffold protein Kidins220, which binds to the TCR, serves to either enhance or reduce TCR signals in iNKT biology, dependent on the developmental stage of the cell. Thus, in the absence of Kidins220, developing thymic iNKT cells receive too strong signals and die by apoptosis, thus limiting the number of iNKT cells. Since iNKT1 cells are more affected than iNKT2 or iNKT17 cells, our data are in line with earlier findings that iNKT1 cells require weak TCR signals to develop (Tuttle et al., 2018). Since Kidins220 binds to many receptors in different cell types (Fiala et al., 2015; Arévalo et al., 2004; Jaudon et al., 2021; Cesca et al., 2011; Scholz-Starke and Cesca, 2016), our new findings might serve as a blueprint to re-examine signal transduction by other receptors.

## Materials and methods

### Mice

Kidins220^+/flox^ mice (Cesca et al., 2012) provided by G. Schiavo (University College London, London, England, UK) were crossed to pLckCre mice (Kidins220lckCre mice). Mice were between 6 and 46 weeks old but for most experiments between 9 and 15 weeks. The C57BL/6Rag2^-/-^ (Rag2 KO), C57BL/6-Ly5.1 (CD45.1) and Kidins220lckCre mice were bred under specific pathogen-free conditions. All mice were maintained in C57BL/6 background. Mice were sex and age matched with litter controls whenever possible. Mice were backcrossed minimum 10 generations to C57BL/6. All animal protocols (G19/151) were performed in accordance with the German animal protection law with authorization from the Veterinär-und Lebensmittelüberwachungsbehörde, Freiburg, Germany.

### Flow cytometry

To gain single cell suspensions from thymus or spleen the organs were mechanically disrupted. Lungs were cut with scissors and treated with 1 mg/ml Collagenase P and 0.1 mg/ml DNAse1 at 37°C for 1 h. Afterwards, the digested lungs were forced through a 70 µm strainer. Single cells are the obtained by gradient centrifugation using percoll. Livers were cut with scissors and forced through a 70 µm strainer. Hepatocytes were gained by centrifugation with percoll supplemented with 100 u/ml heparin. Erythrocytes were lysed using ACK lysis buffer (150 mM NH_4_Cl and 10 mM KHCO_3_) in all single cell suspensions. Afterwards the cells were stained. Flow Cytometry was performed as shown in the company’s instructions using a Gallios (Beckam Coulter) or LSRFortessa (BD Biosciences). Data evaluation was performed using FlowJo X.

### Antibodies and tetramers

The following antibodies were used to stain cells for flow cytometry: PE-labeled anti-CD1d (1B1), PerCP-Cy5.5-labeled anti-CCR7 (4B12), PE-Cy7-labeled anti-CD44 (IM7), PE-labeled anti-CD45.2 (104), FITC Annexin V Apoptosis Detection Kit I, FITC BrdU Flow Kit, AF647-labeled anti-Nur77 (12.14) were purchased from BD Biosciences. APC-labeled anti-CD122 (TM-β1), PB-labeled anti-CD19 (6D5), FITC-labeled anti-CD45.1 (A20), FITC-labeled anti-IFNγ (XMG1.2), PE-Cy7-labeled anti-IL-4 (11B11), PE-labeled anti-MR1 (26.5), FITC-labeled anti-NK1.1 (PK136), PE-Cy7-labeled anti-Sca-1 (D7), PE-labeled anti-Tbet (4B10), FITC-labeled anti-TCRbeta (H57-597), PB-labeled anti-TCRbeta (H57-597) were purchased from Biolegend. PerCP-Cy5.5-labeled anti-CD24 (M1/69), FITC-labeled anti-CD4 (GK1.5), PE-labeled anti-CD4 (RM4-5), PE-Cy7-labeled anti-CD4 (RM4-5), APC-labeled anti-CD44 (IM7), APC-labeled anti-CD5 (53-7.3), PE-Cy7-labeled anti-CD69 (H1.2F3), eFluor660-labeled anti-CD8α (53-6.7), APC-labeled anti-TCRbeta (H57-597) were purchased from eBioscience. PE-Cy7-labeled anti-NK1.1 (PK136) was purchased from Invitrogen. CellEvent Caspase-3/7 green flow cytometry assay kit was purchased from Thermo Fisher Scientific. CD1d tetramers and MR1 tetramers were provided by the NIH Tetramer Core Facility.

### Apoptosis assays

To analyze relative amounts of apoptotic cells, the cells were cultured in RPMI medium supplemented with 10% FBS for 18 h prior to analysis. After staining with surface antibodies, the cells were treated with the Caspase 3/7 reagent (1:200) for 45 min on room temperature (CellEvent Caspase-3/7 green flow cytometry assay kit). This reagent comprises a nucleic acid-binding dye coupled to a peptide (DEVD) which is cleaved by active Caspase 3/7. After cleavage, the dye enters the nucleus and stains nucleic acids enabling the detection of apoptotic cells. For analyzing apoptosis with Annexin V (1:200) it was added for 15 min on room temperature (FITC Annexin V Apoptosis Detection Kit I). For both approaches after incubation the cells were flow cytometrically analyzed without washing. Dead cells were excluded based on FSC and SSC intensities.

### Mixed bone marrow chimera

Bone marrow cells were isolated from CD45.1^+^ mice and mixed in a 1:1 ratio with either isolated bone marrow cells from CD45.2^+^ Ctrl or CD25.2^+^ T-KO mice. 10 million cells of mixed bone marrow cells were intravenously injected into lethally irradiated (9.5 Gy) Rag2 KO mice. Chimeric mice were sacrificed and relative iNKT cell numbers were analyzed 7-8 weeks after injection.

### In vivo BrdU assay

1.5 mg BrdU diluted in 200 µL PBS were i.p. injected into Ctrl and T-KO mice. Mice were sacrificed 16 h or 24 h after injection. Thymocyte proliferation of BrdU-treated mice was assessed following the FITC BrdU Flow Kit (BD) protocol.

### In vivo αGalCer challenge

To test antigen response of iNKT cells in vivo 5 µg of αGalCer (purchased from Biozol) dissolved in 200 µL PBS supplemented with 5.6% sucrose, 0.75% L-histidine and 0.5% Tween20 or only buffer were intraperitoneally injected into Ctrl and T-KO mice. Two hours after injection, mice were sacrificed and splenocytes were isolated to analyze the antigen response of splenic iNKT cells. For this, intracellular cytokines were analyzed as described in the “intracellular staining of cytokines and transcription factors” section.

### Intracellular staining of cytokines and transcription factors

To detect intracellular cytokines and activation markers via flow cytometry, the Cytofix/Cytoperm Kit (BD) was used. Fluorescently labeled CD1dt as well as surface antibodies were stained as usual. Following to this, cells were fixed, permeabilized and stained with anti-IFNγ, anti-IL-4 and anti-Nur77 antibodies in Perm/Wash buffer following the manufacturer’s protocol of the Cytofix/Cytoperm Kit. To stain transcription factors, fluorescently labeled CD1dt and antibodies were used to stain surface molecules. Furthermore, fixation and permeabilization was done according to the manufacturer’s protocol of the Foxp3 / Transcription Factor Staining Buffer Set (eBioscience). Following, cells were incubated with transcription factor antibodies and analyzed by flow cytometry.

## Acknowledgments

We thank Kerstin Fehrenbach and Tabea Reinbold for technical help and Fabrizia Cesca for the Kidins220 floxed mice and discussions. We also thank the Signaling Factory of the Signaling Campus of the University Freiburg. We thank the NIH Tetramer Core Facility (contract number 75N93020D00005) for providing CD1d and MR1 tetramers. We also want to thank BioRender.com for providing schemes of biological systems. This study was supported by the German Research Foundation (DFG) through BIOSS - EXC294 and CIBSS - EXC 2189 (WS and SM,) SFB1381 (A9 to WWS), SFB1160 (B01 to SM) and the DFG grant SCHA-976/7-1 to WWS. KR and GF were supported by the DFG through GSC-4 (Spemann Graduate School).

## Author contributions

LAH, GJF and AMS performed all experiments and data evaluations. The bone marrow chimera experiment was performed with support from KR and SM. BrdU and αGalCer *in vivo* experiments were done with help from RMVC and SM. Apoptosis stains and flow cytometric analysis of iNKT cell stages was done with help from KE, JFH and YT. Data were interpreted by LAH, GJF, AMS and WWS. WWS supervised and conceived the study. LAH and WWS wrote the manuscript. All authors approved the final version of the manuscript.

