## Supplemental figures for "Kidins220 promotes thymic iNKT cell development by reducing TCR signals, but enhances TCR signals in splenic iNKT cells"

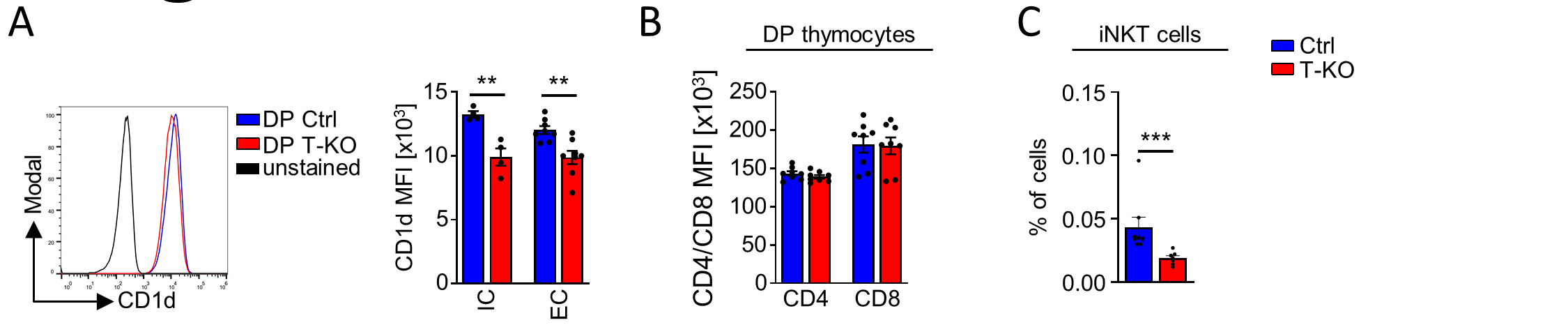

**Figure S1. CD1d, CD4 and CD8 levels on DP thymocytes.**

(A) The CD1d expression level on DP thymocytes was analyzed by intracellular (IC) or surface (EC) stains using anti-CD1d antibodies and flow cytometry. Bar diagrams depict the MFI (n = 4 for IC, n = 8 for EC stains). (B) CD4 and CD8 expression levels on the surface of DP thymocytes were analyzed by stains using anti-CD4 and anti-CD8 antibodies and flow cytometry. (C) Bone marrow cells from CD45.1^+^ WT mice were mixed with bone marrow cells of either CD45.2^+^ Ctrl or CD45.2^+^ T-KO mice in a 1:1 ratio and injected into lethally irradiated Rag2 KO mice. The bar diagram shows relative numbers of CD45.2^+^ Ctrl (blue) and T-KO (red) of thymic iNKT cells (n = 7-8). Statistical analysis for MFIs was performed by two-sided Student’s t test and for relative cell numbers by Mann-Whitney U test; ** p < 0.01; *** p < 0.001. Error bars indicate SEM.

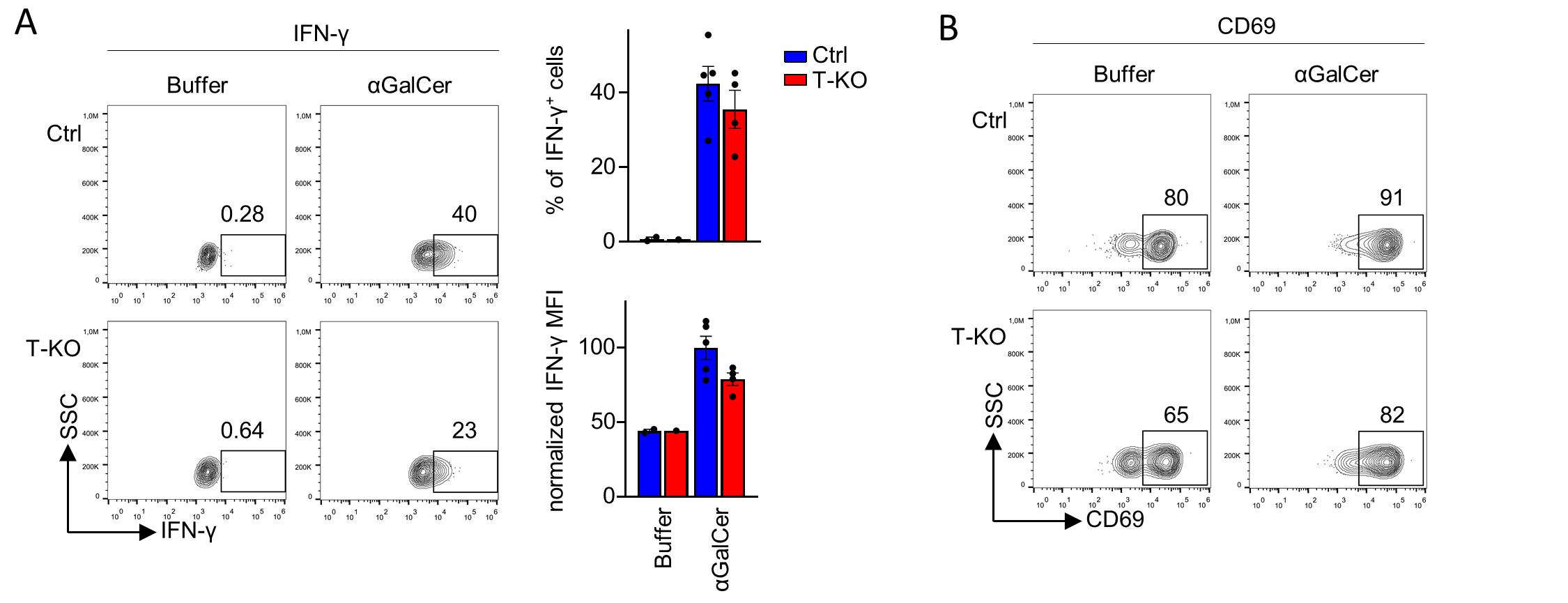

**Figure S2. Splenic T-KO iNKT cells are less responsive to αGalCer.**

Ctrl and T-KO mice were killed 2 h after intraperitoneal injection of αGalCer. Splenocytes were analyzed for (A) IFN-γ^+^ iNKT cells, IFN-γ expression levels as well as for (B) CD69^+^ iNKT cells and CD69 MFI by flow cytometry. Plots were pre-gated on CD1dt^+^ TCRβ^+^ iNKT cells (n = 4-5). Statistical analysis for MFIs was done by two-sided Student’s t test and for relative cell numbers by Mann-Whitney U test. Error bars indicate SEM.

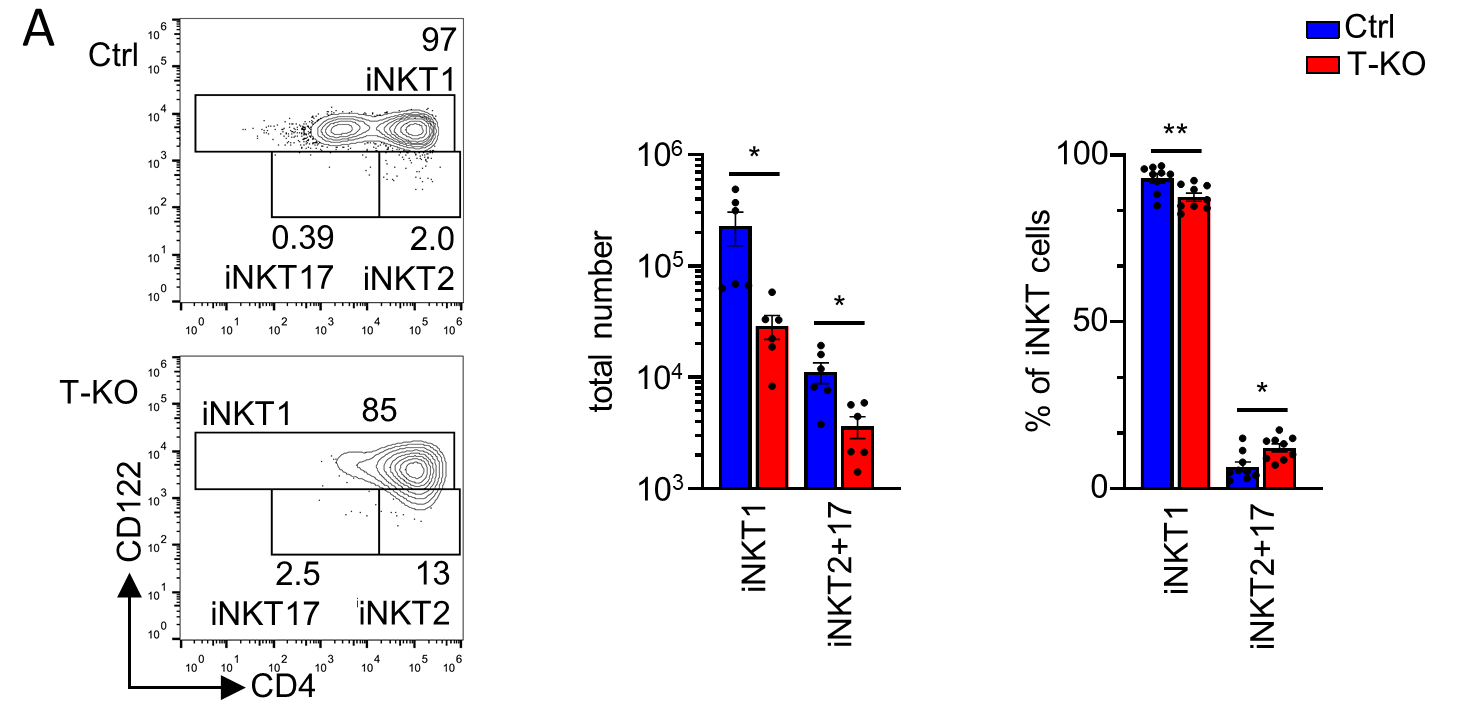

**Figure S3. iNKT1 cell (CD122^+^) numbers are severely reduced in T-KO mice.**

(A) iNKT cell subsets were analyzed using anti-CD122 and anti-CD4 antibodies. iNKT1 cells are CD122^+^ whereas iNKT2 cells and iNKT17 cells are CD122^-^ CD4^+^ and CD122^-^ CD4^‑^, respectively. Samples were pre-gated on CD1dt^+^ TCRβ^+^ iNKT cells. Graphs show cell numbers as well as relative numbers (n = 6-9). Statistical analysis for relative cell numbers by Mann-Whitney U test and for total cell numbers by two-sided Student’s t test; * p < 0.05; ** p < 0.01. Error bars indicate SEM.

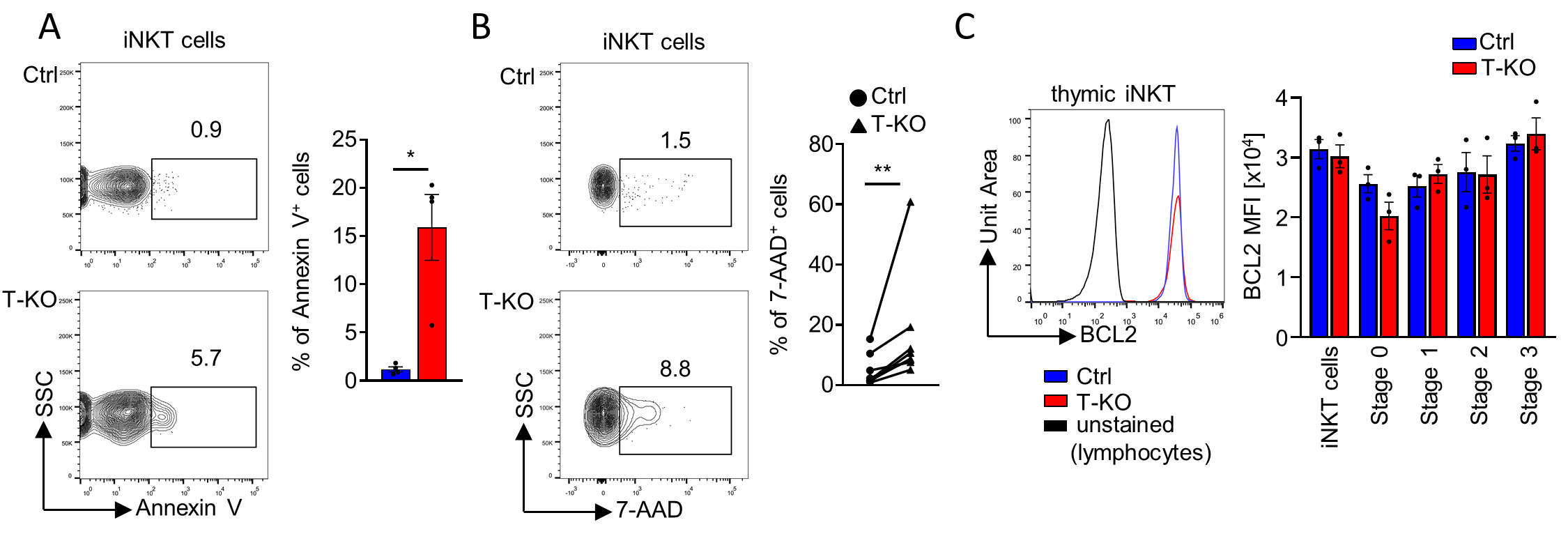

**Figure S4. iNKT cells of T-KO mice show a higher susceptibility to apoptosis than the ones of Ctrl mice.**

(A) For Annexin V-based analysis of apoptosis, thymocytes were cultivated for 18 h in RPMI medium, supplemented with 10% FBS. Thymocytes were stained with Annexin V to detect apoptotic cells and dead cells based on FSC, SSC were excluded. Plots were pre-gated on CD1dt^+^ TCRβ^+^ iNKT cells. The graph depicts the percentage of Annexin V^+^ cells (n = 4). (B) Dead thymocytes were detected using 7-AAD by flow cytometric analysis. Plots were pre-gated on CD1dt^+^ TCRβ^+^ iNKT cells. Graph shows percentage of 7-AAD-positive cells (n = 8). (C) iNKT cells and iNKT cells separated into the different stages (using anti-CD24 antibodies for CD24^-^ stage 0 cells and anti-CD44 and anti-NK1.1 antibodies for stages 1, 2, and 3) were analyzed for BCL2 expression levels using anti-BCL2 antibodies (n = 3). Statistical analysis for relative cell numbers was done by Mann-Whitney U test and by Wilcoxon matched-pairs signed rank test. For MFIs in (C) a two-sided Student’s t test was applied; * p < 0.05; ** p < 0.01. Error bars indicate SEM.
